# Single-cell transcriptomic profiling redefines the origin and specification of early adrenogonadal progenitors

**DOI:** 10.1101/2023.01.09.523195

**Authors:** Yasmine Neirijnck, Pauline Sararols, Françoise Kühne, Chloé Mayère, Serge Nef, Andreas Schedl

## Abstract

Adrenal cortex and gonads represent the two major steroidogenic organs in mammals. Both tissues are considered to share a common developmental origin characterized by the expression of *Nr5a1*/*Sf1*. The precise origin of adrenogonadal progenitors and the processes driving differentiation toward the adrenal or gonadal fate remain, however, elusive. Here we provide a comprehensive single-cell transcriptomic atlas of early mouse adrenogonadal development including 52 cell types belonging to twelve major cell lineages. Trajectory reconstruction reveals that adrenogonadal cells emerge from the lateral plate rather than the intermediate mesoderm. Surprisingly, gonadal and adrenal fates show distinct molecular signatures upon *Nr5a1* induction indicating the two tissues are specified independently. Finally, lineage separation into gonadal and adrenal fate involves canonical versus non-canonical Wnt signaling and differential expression of *Hox* patterning genes. Thus our study provides important insights into the molecular programs of adrenal and gonadal fate choice and will be a valuable resource for further research into early adrenogonadal ontogenesis.

## Introduction

Steroids are essential hormones, regulating a wide variety of developmental and physiological processes including reproduction, blood pressure, and the response to stress and infections. The two main steroidogenic organs in mammals, the adrenal cortex and the gonads (testis and ovary), are thought to share a common developmental origin, the adrenogonadal primordium (AGP). In mice the AGP is specified at around embryonic day (E)9.0-10.0 at the interface of the lateral plate and intermediate mesoderm and is characterized by the expression of the transcription factors *Wt1*, *Gata4*, *Lhx9* and the nuclear steroidogenic factor 1 (*Sf1*, also named *Nr5a1* or *Ad4BP*) (Hu et al., 2013; Ikeda et al., 1994; Wilhelm and Englert, 2002). The AGP is composed of multipotent cells that by E10.5 differentiate into the adrenal primordium (AP) and the gonadal primordium (GP) (Hatano et al., 1996). As the AGP grows and expands, NR5A1^+^ cells delaminate from the coelomic epithelium (CE) and invade the underlying mesenchyme. Ingressing NR5A1^+^ cells along the rostral–caudal axis contribute to the formation of the gonadal primordium. Around E11.5, the bipotential GP initiates its differentiation program according to its chromosomal sex toward testicular or ovarian fate. In contrast the more medial NR5A1^+^ cells located in the rostral region of the AGP migrate to the cranial end of the mesonephros to form the AP. GP and AP cells can be distinguished transcriptionally based on the expression of *Wt1* and *Gata4* that are downregulated in the AP shortly after separation (Bandiera et al., 2013). By E11.5, the AP differentiates as a steroidogenic fetal zone and surrounding mesenchymal cells begin to condense around the primordium that by E12.5-E14.5 will have formed the adrenal capsule (Bandiera et al., 2015).

Despite considerable efforts, the composition and identity of early adrenogonadal progenitors, as well as the transcriptional events that determine their specification and differentiation toward an adrenal or gonadal fate, remain poorly characterized. Furthermore, how the AGP is specified within the mesoderm remains controversial. *Nr5a1* expression analyses and cell lineage tracing studies in chicken have shown that AGP cells originate from the nephrogenic mesenchyme derived from the intermediate mesoderm (IM) (Sekido & Lovell-Badge, 2007; Yoshioka et al., 2005), a concept recently supported by a single-cell transcriptomic study (Estermann et al., 2020a). In contrast, in mammals, *Nr5a1*^+^ AGP cells are first observed in the lateral plate mesoderm (LPM)-derived CE (Hatano et al., 1996; Ikeda et al., 1994), a compartment that contributes to somatic cells of the gonad (DeFalco et al., 2011; Karl and Capel, 1998). However, a recent single-cell transcriptomic analysis of isolated *Osr1*-GFP^+^ cells in mice argues for an IM origin of early adrenogonadal progenitors (Sasaki et al., 2021), highlighting that the ontogeny of the mammalian AGP remains unclear. Here, we generated an unbiased and comprehensive single-cell transcriptomic atlas of early mouse adrenogonadal development from E9.0 to E12.5. This atlas allowed us to characterize the progenitor cells present in the AGP, map their ontogeny and characterize the lineage specification and differentiation towards adrenal and gonadal primordia.

## Results

### A single-cell transcriptomic atlas of early adrenogonadal development

To generate a gene expression atlas of early adrenogonadal development, we performed single-cell transcriptome profiling using a high throughput droplet-based system (10X Genomics). Single cells were isolated at the time of AGP specification (E9.0), AGP separation (E10.5), gonadal sex determination (E11.5) and subsequent differentiation (E12.5) (Bullejos and Koopman, 2001; Ikeda et al., 1994). Entire posterior trunks (E9.0), urogenital ridges (E10.5 and E11.5), and gonads (E12.5) were isolated from XY and XX *Nr5a1^Tg^* mouse embryos. Adrenals glands (E12.5) were harvested only from XY *Nr5a1^Tg^* embryos. Our dataset includes a total of 72,273 cells and 18,913 genes, with a median of 20,260 unique transcripts (UMIs) per cell, 4,558 genes per cell and 4.5 UMIs/gene per cell (**Fig. S1A,B**). Visualization of the single-cell transcriptomes in a uniform manifold approximation and projection space (UMAP) (Becht et al., 2019)) shows that while male and female cells overlap at early stages, a subset of male cells emerges at E11.5, concomitant with gonadal sex determination (Bullejos and Koopman, 2001). By E12.5, the cells comprising the developing ovaries, testes and adrenal glands are clearly separated, reflecting diversification of the adrenogonadal cell types (**Fig. 1A**). We applied unsupervised clustering (see methods) to classify cells based on transcriptomic similarities, resulting in 52 clusters (**Fig. 1B**). Differential expression analysis identified specific or enriched genes in each cluster that were subsequently cross-referenced with previously established markers (**Table S1**). Cell types were assigned manually using *a priori* knowledge, combining expression of known specific/enriched marker genes, developmental age, sex and sample origin of the cells (**Fig. 1B, C**, **Fig. S1C,D**, **Fig. S2** and **Table S1**). We mapped the 52 clusters (hereafter abbreviated as C) into 12 major lineages (**Fig. 1C** and **Fig. S2**): hematopoietic/endothelial, germ cells, neural/sympathoadrenal, notochord, paraxial mesoderm, nephrogenic, mesonephros-associated cell types, gonadal steroidogenic, adrenal steroidogenic, lateral plate mesoderm, adrenogonadal progenitors, and supporting cell lineages.

**Figure 1.**
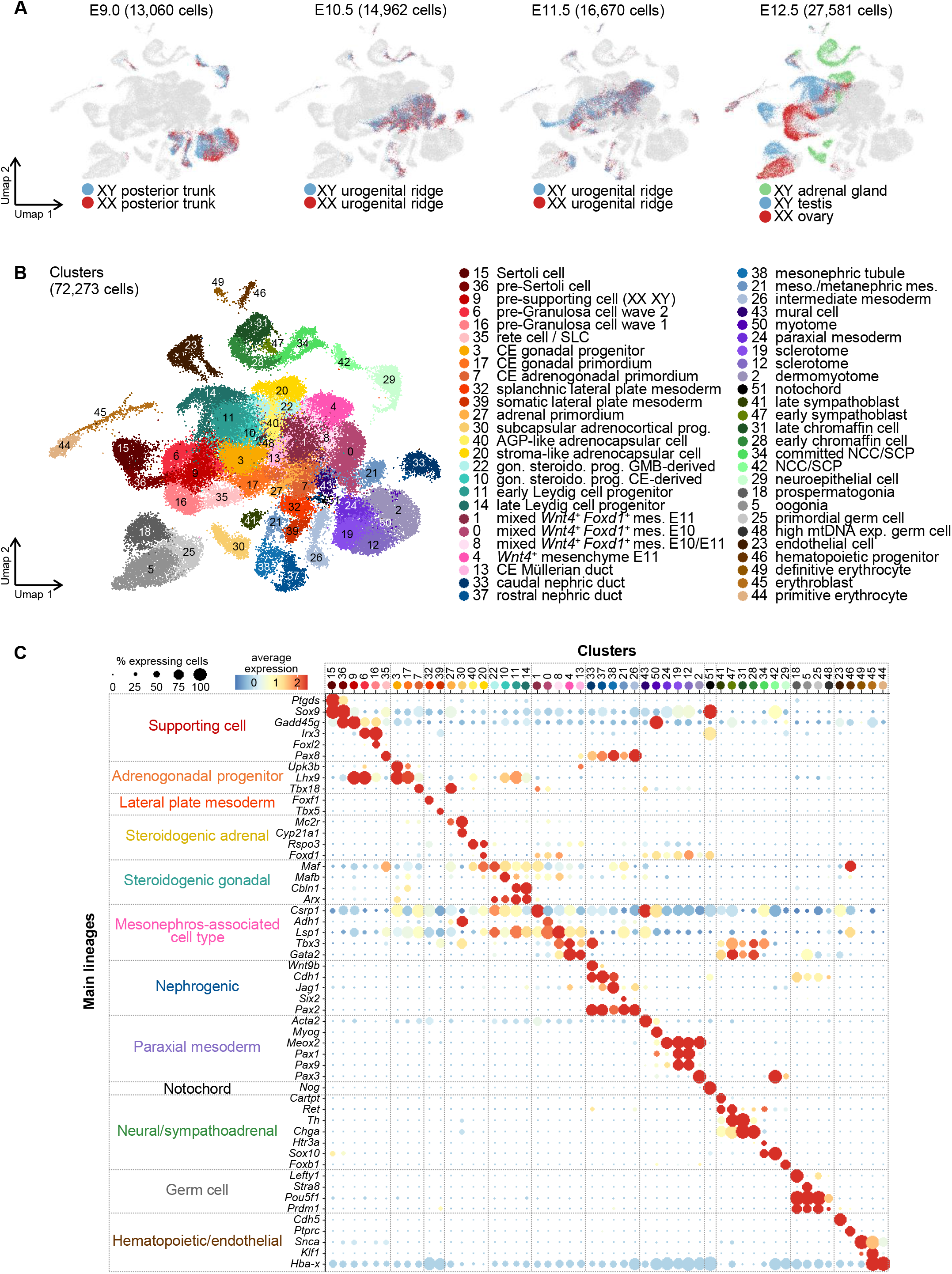
A single-cell transcriptomic atlas of early adrenogonadal development. **A, B.** UMAP visualization of the 72,273 transcriptomes colored by age, sex, sample type (A), and by clusters (B). Cell type annotation is shown on the right. Abbreviations: AGP, adrenogonadal primordium; CE, coelomic epithelium; GMB, gonad/mesonephros border; gon. steroido. prog., gonadal steroidogenic progenitor; mes., mesenchyme; meson., mesonephric; mtDNA exp., mitochondrial DNA expressing; NCC, neural crest cell; SLC, Supporting-like cell; SCP, Schwann cell precursor; term. diff., terminally differentiated. **C.** Dot plot showing expression of selected marker gene (y axis) per cluster (x axis). The size of the dot represents the percentage of cells in the cluster expressing the gene and the color indicates the level of expression (log normalized counts). Major cell lineages are displayed on the left. (See also figures S1-S2 and note S1).

Adrenogonadal progenitors were identified by expression of early adrenogonadal markers (*Gata4*, *Lhx9*, *Nr5a1*) together with coelomic epithelium markers (*Upk3b* (Rudat et al., 2014), *Myrf* (Hamanaka et al., 2019)). It is composed of cells of both sexes from E10.5 (C17) and from E11.5/E12.5 (C3), annotated as “coelomic epithelium gonadal primordium” and “coelomic epithelium gonadal progenitor”, respectively. The fate of CE progenitors have been shown to differ before and after E11.2, preferentially giving rise to supporting and steroidogenic cells, respectively (DeFalco et al., 2011; Karl and Capel, 1998). The fact that early and late CE cells in our dataset are transcriptionally distinct, and that CE cells at E11.5/E12.5 cluster close to the steroidogenic lineage, is in agreement with this work. An additional population of *Gata4*^+^/*Lhx9^low^*/*Upk3b^low^*/*Myrf*^+^ cells is present at E10.5 (C7). *Nr5a1*, whose expression is known to be downstream of *Gata4* and *Lhx9*, is scarcely detected in this cluster (Birk et al., 2000; Hu et al., 2013; Wilhelm and Englert, 2002). C7 thus corresponds to newly specified adrenogonadal cells and is annotated “coelomic epithelium adrenogonadal primordium”.

The supporting cell lineage comprises XY *Sox9*^+^/*Ptgds^low^* pre-Sertoli and *Sox9*^+^/*Ptgds^high^* Sertoli cells (C36 and C15, respectively), *Gadd45g*^+^ pre-supporting cells originating from both sexes at E11.5 (C9), and two clusters consisting of E12.5 ovary cells expressing the pre-Granulosa cell markers *Irx3* and *Fst* (C6 and C16). While C16 preferentially expresses *Foxl2*/*Hmgcs2*/*Akr1cl*, C6 expresses higher amounts of *Lhx9*/*Gng13*, which correspond to the first and second wave of pre-Granulosa cells recently described in (Niu and Spradling, 2020). C35 contains cells of both sexes at E11.5/E12.5, co-express *Nr5a1*, *Wt1*, *Sox9* as well as *Pax8*, and is therefore referred to as “rete cell/supporting-like cell (SLC)”(Kulibin and Malolina, 2020; Omotehara et al., 2020), at the origin of the rete testis and rete ovarii (Mayère et al., 2021a).

Progenitors of the gonadal steroidogenic lineage were identified by *Arx* expression (Miyabayashi et al., 2013). *Insl3* C11 and *Insl3*^+^ C14 were exclusively composed of testicular cells at E12.5 and were therefore referred to as early and late Leydig cell progenitors, respectively. C10 and C22 on the other hand, were derived from both sexes and express different levels of *Lhx9*, *Mafb* and *Maf* (DeFalco et al., 2011). *Lhx9*^+^/*Mafb^high^*/*Maf^low^* C10 is therefore annotated as “gonadal steroidogenic progenitor coelomic epithelium-derived”, whereas *Lhx9*^-^/*Mafb^low^*/*Maf^high^* C22 is referred to as “gonadal steroidogenic progenitor gonad/mesonephros border-derived”.

The adrenal steroidogenic lineage contains *Nr5a1*^+^ steroidogenic cells at E12.5 that co-express adrenal-specific genes (*Mc2r*, *Cyp21a1*) and adrenal steroidogenic progenitor markers (*Wnt4*, *Shh^low^*, *Gli1^low^*); thus, they have been named “subcapsular adrenocortical progenitor” (C30) (King et al., 2009). Adrenal gland samples at E12.5 also contain two *Nr5a1^-^* /*Gli1^+^*/*Ptch1^+^* non-steroidogenic populations that could likely represents SHH-responsive capsular steroidogenic progenitors (C20 and C40) (King et al., 2009). While C40 displays an early adrenogonadal transcriptomic signature (*Lhx9*, *Gata4*, *Nr0b1*, *Tbx18*), C20 cells express higher levels of capsular genes *Tcf21* and *Wt1* (Bandiera et al., 2013; Wood et al., 2013) and stromal genes (*Foxd1*, *Dcn*, *Olfml3*). C40 and C20 were therefore annotated as “adrenogonadal-like adrenocapsular cells” and “stroma-like adrenocapsular cells”, respectively. The adrenal primordium is identified at E10.5 by *Nr5a1*^+^ cells that differs from the gonadal primordium by low expression level of *Wt1*/*Gata4*/*Nr0b1* (C27) (Bandiera et al., 2013; Zubair et al., 2008). This cluster expresses *Mc2r*, *Star* and *Mgarp*, indicating that a steroidogenic transcriptional program is already initiated at this early stage. An exhaustive description of additional lineages/cluster is available in **note S1**.

Overall, our dataset provides a comprehensive overview of the expected adrenogonadal cell types as well as surrounding tissues, present before and during AGP specification, as well as the more differentiated adrenogonadal lineages. This adrenogonadal developmental atlas is freely accessible through an interactive web portal (http://lmedapp731.unige.ch:5006/) allowing to query for genes of interest per cell types, age and sex.

### *In silico* reconstruction of adrenogonadal primordium separation refines known and novel marker genes kinetics

To investigate the transcriptional programs underlying AGP specification and fate choice toward adrenal or gonadal identity, we focused the analysis on cells annotated as “coelomic epithelium adrenogonadal primordium” (C7), “coelomic epithelium gonadal primordium” (C17) and “adrenal primordium” (C27). These cells were visualized using a force-directed layout resulting in a V-shape structure with AGP cells at the base of two branches composed of cells from the gonadal or adrenal primordia reflecting AGP separation (**Fig. 2A**). A refined clustering revealed high heterogeneity and transitional cellular states with 8 subclusters (subC) (**Fig. 2B**). Gene expression analysis together with partition-based graph abstraction (PAGA) (Wolf et al., 2019) defined a common trajectory made of “early AGP” (subC1), “late AGP” and “dividing late AGP” cells (subC1, 6 and 3 respectively), a gonadal trajectory made of “early” and “late gonadal primordium” cells (subC4 and 0 respectively) and an adrenal trajectory made of “early” and “late adrenal primordium” cells (subC2 and 5 respectively) (**Fig. 2C-E** and **Table S2**). We have recently shown that rete progenitor cells or “SLC” are specified as early as E10.5 and represent a coelomic epithelium-derived lineage diverging from the gonadal somatic cell lineage (Mayère et al., 2021a). Therefore, the “early rete cell/SLC” cluster (subC7), which represents the earliest cells of the SLC lineage, was not taken into account for further analysis. Expression of AGP markers along pseudotime shows that *Gata4*/*Wt1* detection precedes *Lhx9*/*Emx2* expression, followed by *Nr5a1* induction (**Fig. 2E**) in accordance with previous reports (Hu et al., 2013; Wilhelm and Englert, 2002). Shortly after *Nr5a1* detection, these genes adopt gonadal- and adrenal-lineage specific expression profiles: they are downregulated in cells of the adrenal lineage concomitant with *Mc2r*/*Nr5a1* upregulation, whereas they are further upregulated in gonadal lineage together with induction of the supporting and steroidogenic cell markers *Gadd45g* and *Cbln1*. Interestingly, the *Nr5a1*^+^ cellular state preceding lineage diversification (highlighted in two dotted red lines in **Fig. 2E**) is composed in majority by cells of cluster 2 (“early AP”) and cluster 4 (“early GP”) of the adrenal and gonadal lineages, respectively. This indicates that *Nr5a1* expression is induced in two distinct cell populations. Taken together, pseudotime ordering recapitulates the sequential expression kinetics and dynamics of known AGP markers, providing a robust tool to investigate the global transcriptomic signatures associated with each differentiation steps. Importantly, it challenges the view of a common *Nr5a1*^+^ progenitor and suggests that CE cells are already specified toward either adrenal or gonadal fate before delamination and ingression within the underlying mesenchyme.

**Figure 2.**
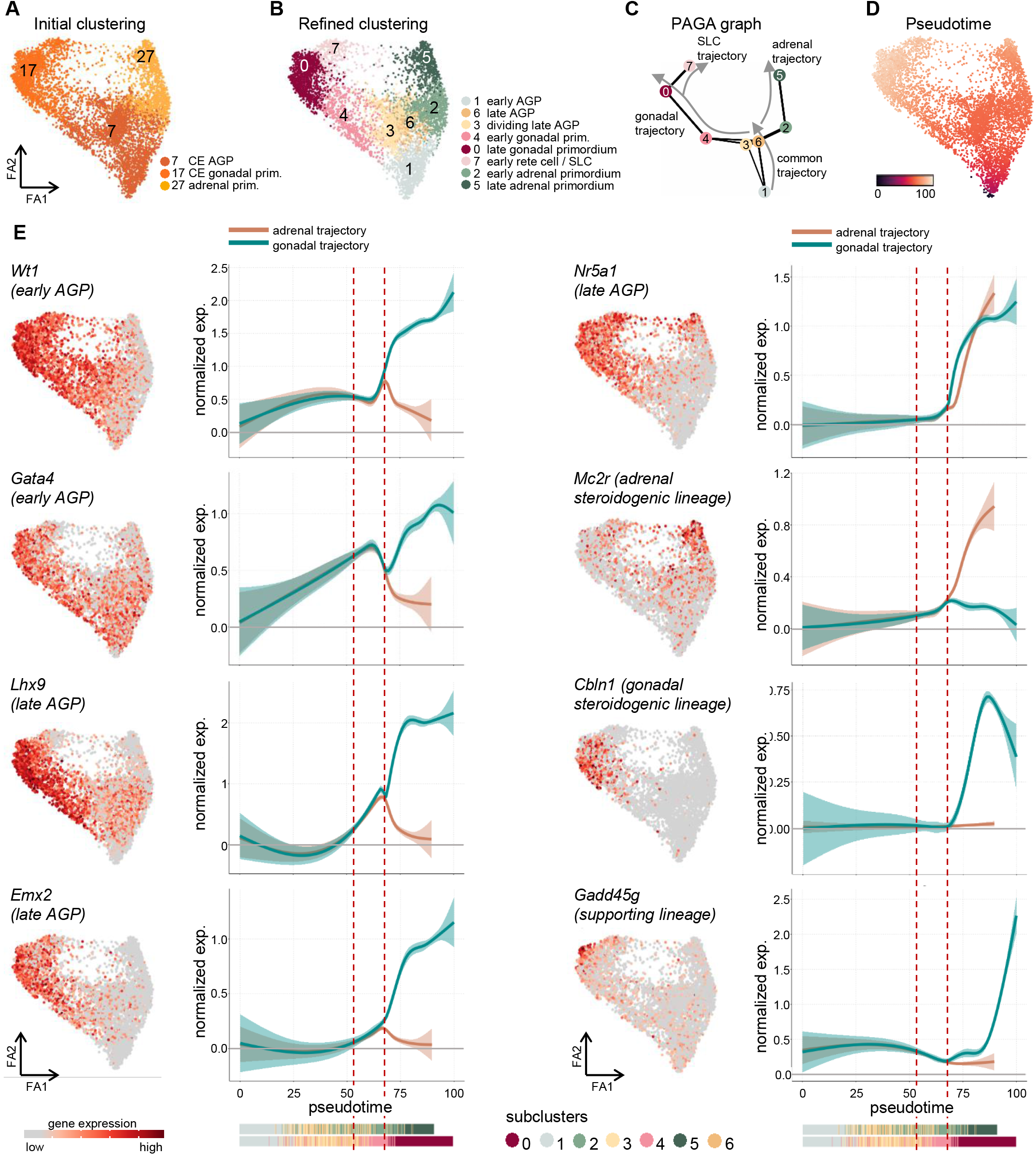
*In silico* reconstruction of adrenogonadal primordium separation refines the kinetics of known and novel marker genes. **A,B.** Visualization of the 5,128 transcriptomes in a force-directed layout colored by initial clustering (A) and new refined clustering. Cell type annotation is shown on the right. Abbreviations: AGP, adrenogonadal primordium; CE, coelomic epithelium; SLC, Supporting-like cell. **C.** PAGA representation of the refined clustering. Each node is a cluster and the edges between clusters represent connectivities. Edges with low connectivities (below 0.3) are not represented. Arrows indicate the common, gonadal, adrenal and SLC trajectories. **D.** Visualization of the transcriptomes colored by pseudotime. **E.** Expression levels of representative marker genes in a force-directed layout (left) and along pseudotime (right, y axis: normalized expression, x axis: pseudotime). The two vertical dotted red lines indicate *Nr5a1*^+^ cellular state preceding lineage diversification.

### Adrenal fate choice is associated with inhibition of canonical Wnt signaling and enhanced Rho signaling gene expression

To characterize the transcriptomic programs underlying adrenal versus gonadal cell commitment, we selected 1,737 genes with dynamic expression patterns along pseudotime (see methods) and grouped them according to their expression similarities. We obtained 22 gene profiles (P) (**Table S3**) including 10 and 7 profiles for which gene expression increased during gonadal and adrenal specification, respectively, and 5 profiles displaying sequential and transitory expression dynamics common to both lineages (**Fig. 3**). Genes of P16, 13 and 7 are upregulated during gonadal progression. Gene ontology (GO) analysis (**Table S4** and **Fig. 3**) associated these genes with terms such as “regulation of reproductive process” (P16, *Sulf1*, *Mafb*) “positive regulation of gonad development” (P13, *Wt1*, *Amhr2*) and “reproductive structure development” (P7, *Nr0b1*, *Lef1*, *Igf1*, *Lhx9*, *Rspo1*, *Emx2*, *Msx1*, *Lgals7*, *Lgr4*). One profile contains gonadal-specific CE genes (P20, *Upk3b*, *Myrf*) together with genes coding for keratins, indicating CE maturation accompanies gonadal specification.

**Figure 3.**
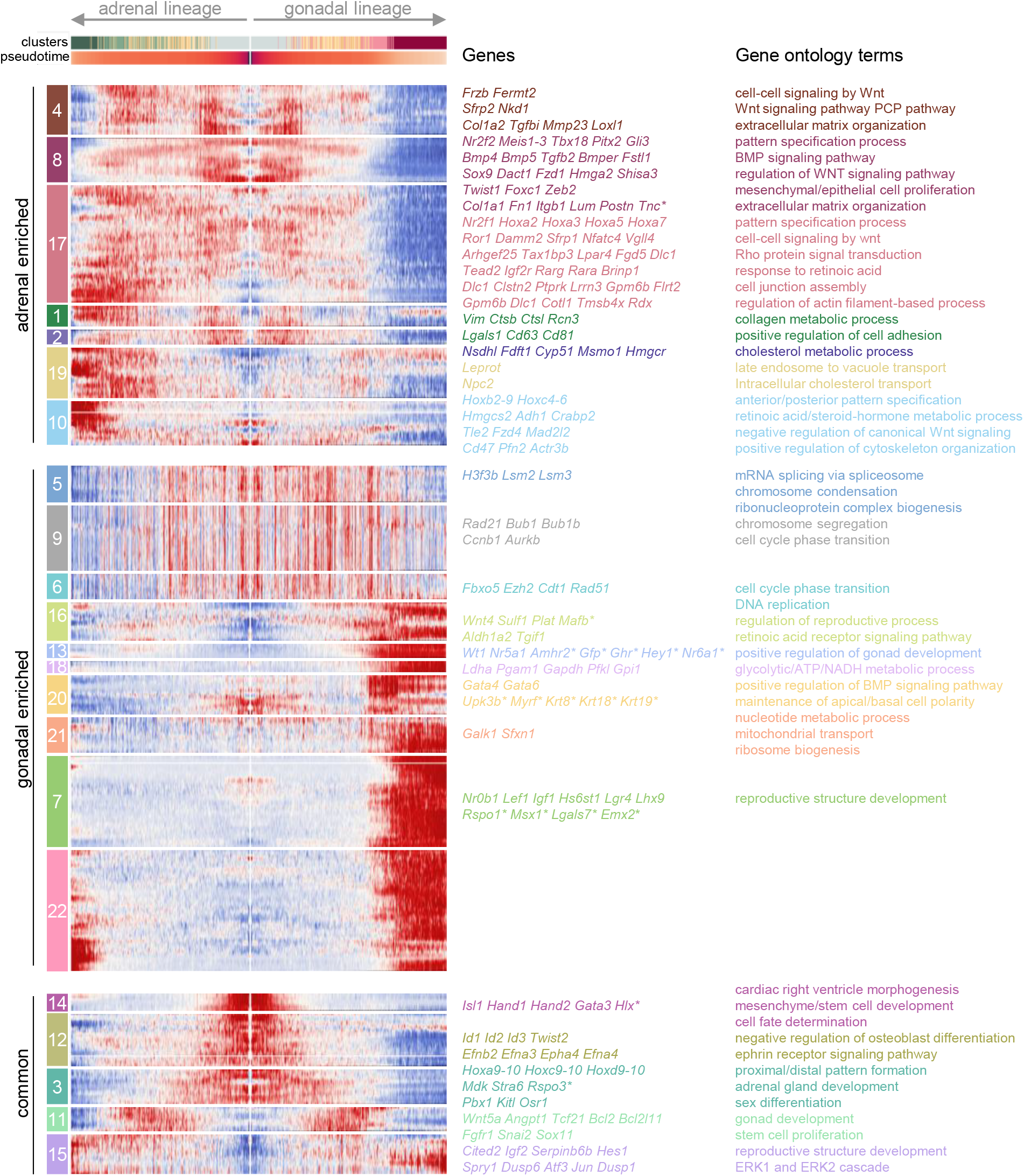
Common and divergent transcriptional programs during adrenal and gonadal fate specification. Mirror heatmap showing the expression profiles of 1,737 genes along pseudotime. Cells were ordered according to their pseudotime score with early AGP cells in the center of the figure (score 0) and endpoints of the adrenal and gonadal lineage on the left and right sides, respectively. The 22 gene profiles are indicated on the left, and their associated genes and GO terms are shown on the right. (See also figure S3).

Regarding adrenal enriched programs, genes of P4, 8, 17, 1 and 2 are expressed in both lineages prior adrenal or gonadal specification, are progressively downregulated in gonadal cells whereas expression is maintained or further increased in adrenal cells. GO analysis associated these genes with “cell-cell signaling by Wnt”, “Wnt signaling pathway PCP pathway” and “regulation of WNT signaling pathway”. Genes belonging to these terms (*Sfrp1*, *Sfrp2*, *Nkd1*, *Sox9*, *Dact1*, *Fzd1*, *Hmga2*, *Shisa3*, *Ror1*, *Daam2*, *Nfatc4*, *Vgll4*, **Fig. S3A**) are all annotated as “negative regulation of canonical Wnt signaling pathway” in the GO database. In line with their opposite expression levels between adrenal and gonadal lineages, the target of the canonical Wnt signaling *Lef1* (P7) is induced in gonadal but not adrenal cells. Adrenal programs involve genes associated with “Rho protein signal transduction” (P17), “regulation of actin filament-based process” (P17), “positive regulation of cytoskeleton organization” (P10), “extracellular matrix organization” (P4 and 8) and “collagen metabolic process” (P1), suggesting non-canonical Wnt activation of Rho signaling is at play in AGP and AP cells to positively modulate cell movement and interaction with extracellular matrix. This is consistent with the morphogenetic events taking place in the AGP, as cells delaminate from the CE and undergo an epithelial-to-mesenchymal transition (EMT). Accordingly, the EMT inducers *Twist1*, *Zeb2* and *Foxc1* (Meyer-Schaller et al., 2019) are regulated in a similar manner (P8).

### Early AGP gene signatures indicate a lateral plate mesoderm origin and dynamic *Hox* expression programs

To get insights into the gene programs underlying early AGP cells specification prior *Nr5a1* expression, we focused on the gene profiles displaying high expression at the start of the pseudotime. Among the 22 profiles, P14 included genes that have an early and robust high expression levels and then are rapidly downregulated as the cells progress to the late AGP state (**Fig. 3**). These genes are associated with the GO term “cardiac right ventricle morphogenesis” including *Isl1*, *Hand1*, *Hand2*, *Gata3*, which are known markers of the proepicardium, a CE-derived structure (**Fig. S3B**). P14 also includes *Hlx* expressed in the septum transversum (Lints et al., 1996). The finding that the early AGP gene signature reflects that found in proepicardium/septum transversum cells, which are derived from the LPM (Cano et al., 2016), suggests that the AGP also originates from the LPM rather than the IM. To confirm this hypothesis, we compared the PAGA connectivity’s scores, indicating potential ontogenic relationships, among mesodermal-derived cell types of our whole dataset (E9.0 to E12.5). Indeed, this analysis showed that AGP cells (C7 of our initial clustering) are transcriptionally closer to splanchnic LPM at E9.0 (C39) than IM at E9.0 and meso/metanephric mesenchymal cells at E10.5 (C26 and C21, respectively) (**Fig. 4A**).

**Figure 4.**
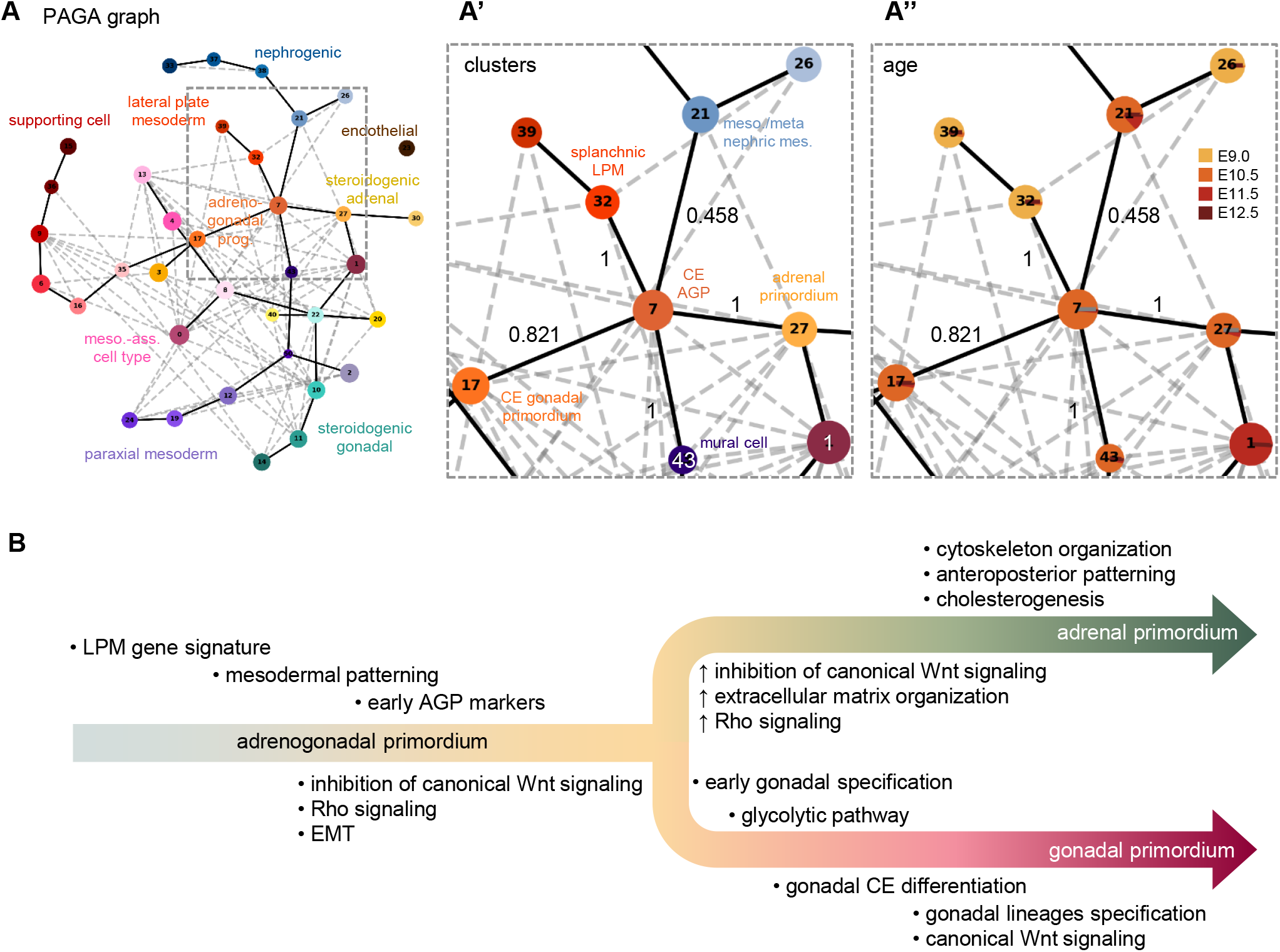
Lineage reconstruction of mesodermal-derived cell types from E9.0 to E12.5. **A.** PAGA representation of the mesodermal-derived cell clusters from figure 1B. Each node is a cluster and the edges between clusters represent connectivities. Edges with low connectivities (below 0.25) are not represented. Major lineages are indicated. The inset shows the connectivity scores (black numbers) of cluster 7 (CE AGP) with clusters 21, 27, 32 and 43 and are colored by clustering (**A’**) and by age (**A”**). **B.** Model of cell lineage specification during adrenogonadal primordium separation.

The other profiles with high gene expression in early AGP cells include P12 and P3 which are downregulated during both adrenal and gonadal cells specification, and P4, P8 and P17 which are downregulated in the gonadal lineage only. GO annotations common to these profiles include “proximal/distal pattern formation” and “pattern specification process” with corresponding genes including several *Hox* genes (P3 and 17) and mesodermal markers such as *Nr2f1*, *Nr2f2*, *Meis1-3*, *Pitx2* and *Gli3* (P8 and 17). *Hox* genes encode transcriptional regulatory proteins and are organized in four genomic clusters (*Hoxa, b, c* and *d*) expressed from 3’ to 5’ along the antero-posterior body plan (Deschamps and Duboule, 2017). Interestingly, *Hoxa9-10, Hoxc9-10, Hoxd10* genes are downregulated in both lineage (P3) while the expression of *Hoxa2,3,5,7* genes is maintained in adrenal cells (P17). In addition, *Hoxb2-9* and *Hoxc4-6* expression increased as cells adopt an adrenal fate (P10) (**Fig. S3C**). Several *Hox* genes have been shown to be abundantly expressed in AP cells where they directly bind and positively regulate *Nr5a1* expression (Zubair et al., 2006). Since our analysis shows differential expression of the *Hox* genes along adrenal and gonadal fate commitment, and since the AP is specified in the more anterior region of the genital ridge, it is possible that specific sets of *Hox* genes determine adrenal versus gonadal fate choice.

## Discussion

Although adrenogonadal development has been studied for several decades, the composition and molecular signatures of early adrenogonadal progenitors is still incomplete. In addition, the genetic programs and regulatory factors orchestrating the separation of the AGP into two distinct primordia remain poorly characterized. Here we generated a comprehensive single-cell transcriptomic atlas of mouse adrenogonadal cells and surrounding tissues collected before, during and after AGP separation, without prior selection of cells by lineage markers. This provided us with in a detailed atlas comprising 52 cell types, which is a valuable resource for the adrenogonadal research community. These data are freely available to the scientific community in an interactive website (http://lmedapp731.unige.ch:5006/).

Using this atlas we showed that adrenogonadal primordium and the gonadal and adrenal primordia at E10.5 display distinct transcriptomic identities. Lineage reconstruction and pseudotime analysis, allowed us to uncover the dynamic genetic programs at play during AGP cell specification toward the adrenal or gonadal fate, unraveling differential cellular processes underlying this fate choice (model depicted in **Fig. 4B**). We found that the canonical Wnt pathway is actively repressed in uncommitted AGP cells and this repression is maintained or further increased as the cells progress through the adrenal lineage. In contrast, cells adopting a gonadal fate release this inhibition and activate the canonical Wnt pathway required for their proliferation (Chassot et al., 2012). On the other hand, cells of the AGP and the adrenal primordia appear to activate the planar cell polarity pathway as they undergo Rho signaling-dependent rearrangement of actin filament and cytoskeleton, concomitantly with active interaction with extracellular matrix components and epithelial-to-mesenchymal transition. These processes may be important for the cells with adrenal fate to move to a more dorso-medial position than the gonadal ridge, allowing proper positioning of the future fetal cortex within the urogenital tract. Indeed, between E11.5-E14.5, the fetal adrenal undergoes extensive morphogenetic events, such as invasion by Schwann cell precursors to form the adrenal medulla, and encapsulation by surrounding mesenchymal cells to form the adult adrenal cortex (Bandiera et al., 2015). Furthermore, the dorso-medial positioning of the fetal cortex may be important to reach an “adrenal niche” presenting optimal cues from surrounding tissues, for proper differentiation into fetal adrenal steroidogenic cells. Interestingly, mice mutant for *Wnt4*, a gene that has been reported to induce non-canonical Wnt signaling in several organs (Eliazer et al., 2019; Louis et al., 2008; Tanigawa et al., 2011), exhibit mislocalization of adrenal cells at the anterior pole of the gonads (Heikkilä et al., 2002; Jeays-Ward et al., 2003; Val et al., 2006), supporting our model that non-canonical adrenal-specific Wnt signaling allows for proper separation of the gonadal and adrenal primordia. Considering the differential combinations of *Hox* genes expressed in cells destined to become adrenal and gonadal cells, it appears that the specification/separation of adrenal and gonadal primordia involves a complex integration of positional information as well as dedicated cellular processes allowing proper patterning of adrenogonadal ridges.

Careful analysis of the expression profile of the known early adrenogonadal markers along pseudotime revealed that *Nr5a1* is expressed relatively late within the AGP differentiation window, in cells which are already committed toward gonadal or the adrenal fate. This reveals that the use of *Nr5a1* as a marker of the CE to define a common AGP is incorrect and raises the question whether CE cells are already specified toward a given fate before delamination and ingression within the underlying mesenchyme. The expression profile of *Sfrp2*, a negative regulator of the canonical Wnt pathway, and of the genes encoding the neurofilament proteins *Nefl* and *Nefm*, appear to be good candidates to test this hypothesis, as we found them to be expressed early during adrenal pseudotime. Of note, our conclusions are in accordance with a recent study in human and monkey which revealed adrenal and gonadal cells emergence within the coelomic epithelium is spatially distinct, in the “adrenogenic” and “gonadogenic” CE, respectively (Cheng et al., 2022). Therefore, distinct specification of these two lineages appears to be common to both mouse and human.

The origin of early adrenogonadal cells is disputed, as it remains unclear whether they are derived from the lateral plate or the intermediate mesoderm. Indeed, the only cell lineage tracing studies of coelomic epithelial cells in mouse were performed after AGP separation (DeFalco et al., 2011; Karl and Capel, 1998). We used our single-cell dataset comprising cells from all three layers of mesodermal derivatives (paraxial, lateral and intermediate) present at E9.0 in the posterior trunk to show that in mouse, early AGP cells are ontogenically related to the splanchnic lateral plate mesoderm. The relevance of different origins for the AGP among vertebrates-LPM in mouse (this study) and IM in chicken (Estermann et al., 2020a; Sekido and Lovell-Badge, 2007; Yoshioka et al., 2005)-is not clear, but it has been proposed to be linked to differences in mesonephros excretory function and/or evolution of master sex-determining switches genes (Estermann et al., 2020b). Interestingly, we observed discrepancies between *Nr5a1*-*eGFP* transgene and endogenous *Nr5a1* expression, with *GFP* being expressed in a subset of *Pax2*^+^/*Nr5a1*^-^ intermediate mesoderm at E9.0 (data not shown). Given that the *Nr5a1*-*eGFP* transgene and the endogenous *Nr5a1* locus differs in their regulatory regions (for comparison see (Stallings et al., 2002; Zubair et al., 2006)), the observed induction of *Nr5a1*-*eGFP* in mouse IM could reflect an ancestral adrenogonadal fate specification competence which has been switched off during evolution. Comparative mouse and chicken mesodermal single-cell transcriptomic and epigenomic profiling, could provide insights into the evolutionary divergence of regulatory processes of adrenogonadal specification.

In summary, the here described comprehensive atlas of adrenogonadal development allowed us to determine the precise origin of the adrenal cortex and gonads in the mouse. Our findings that development of these two organs diverges before the induction of *Nr5a1* indicates that this gene should no longer be used as a marker for the common primordium.

## Acknowledgments

We thank the teams of the Flow Cytometry Facility (Faculty of Medicine, University of Geneva), the Animal Facility (Faculty of Medicine, University of Geneva), the DNA sequencing platform (Health 2030 genome center, Geneva) and Dr Christelle Borel (GEDEV department, University of Geneva) for help with the 10X Chromium technology.

This work was supported by the Fondation pour la Recherche Médicale (grant number ARF201909009270 to Y.N.), the Swiss National Science Foundation (grant numbers 31003A_173070 and 310030_200316 to S.N.), the Département de l’Instruction Publique of the State of Geneva (to S.N. and Y.N), La Ligue Contre le Cancer (Equipe Labelisée to A.S.) and the ANR (ANR-11-LABX-0028-01 and ANR-18-CE14-0012 to A.S.).

## Author contributions

Conceptualization, Y.N., S.N. and A.S.; Methodology, Y.N. and P.S.; Software, P.S. and C.M.; Formal analysis and Investigation, Y.N., P.S., F.K. and C.M; Resources, S.N.; Data Curation, Y.N., P.S. and C.M.; Writing – Original Draft, Y.N.; Writing – Review & Editing, Y.N., P.S., S.N. and A.S.; Supervision, Y.N., S.N. and A.S.; Funding Acquisition, Y.N., S.N. and A.S.

## Methods

### Mouse posterior trunks, urogenital ridges, adrenal glands, testes and ovaries collection

All animal work was conducted according to the ethical guidelines of the Direction Générale de la Santé of the Canton de Genève (experimentation ID GE/57/18). *Nr5a1*-*eGFP^Tg^* mouse strain was described previously (Stallings et al., 2002) and has been maintained on a CD1 background. E9.0 (18±3 somites), E10.5 (8±2 tail somites (ts)), E11.5 (19±4 ts) and E12.5 embryos from timed matings (day of vaginal plug=0.5) were collected and the presence of the *Nr5a1*-*eGFP* transgene was assessed under UV light. Sexing of E9.0, E10.5 and E11.5 embryos was performed by PCR with a modified protocol from (McFarlane et al., 2013). Each condition (organ, sex and developmental stage) was performed in two independent replicates.

### Single cell suspension and library preparation

Posterior trunks, urogenital ridges and adrenal glands were enzymatically dissociated at 37°C between 30 and 40 minutes using the Papain dissociation system (Worthington LK003150). Cells were resuspended in DMEM (Gibco 11885084) supplemented with 2% fetal bovine serum (Sigma-Aldrich F2442), filtered through a 70 μm cell strainer and stained with Draq7 (Beckman Coulter B25595). Viable single cells were collected on a BD FACSAria II by excluding debris (side scatter *vs*. forward scatter), dead cells (side scatter *vs*. Draq7 staining), and doublets (height *vs*. width). Testes and ovaries were enzymatically dissociated 10 minutes at 37°C using Trypsin-EDTA 0.05% (Gibco 25300054). After centrifugation and counting, 3000 to 7000 single cells were loaded on a 10x Chromium Controller (10x Genomics). Single-cell RNA-Seq libraries were prepared using the Chromium Single Cell 3’ Reagent Kit v2 (10x Genomics PN-120237 PN-120236 PN-120262) according to manufacturer’s protocol.

### Sequencing

Library quantification was performed using the Qubit fluorometric assay with dsDNA HS Assay Kit (Invitrogen Q32851). Library quality assessment was performed using a Bioanalyzer Agilent 2100 with a High Sensitivity DNA chip (Agilent 5067-4626). Libraries were diluted, pooled and sequenced on an Illumina HiSeq4000 using a paired-end, single index sequencing mode as follow: 26 base-pair (bp) Read1, 98 bp Read2 and 8 bp i7 index. Libraries were sequenced at a targeted depth of 100,000 to 150,000 reads per cell. Sequencing was performed at the Health 2030 Genome Center, Geneva.

### Single-cell RNA sequencing analysis

Computations were performed at the Vital-IT Centre for high-performance computing of the Swiss Institute of Bioinformatics (http://www.vital-it.ch).

Demultiplexing, alignment and UMIs quantification were performed with the cellranger software suite (version 2.1, 10X Genomics). The mouse reference genome GRCm38.p5 and the Gencode annotation (release M15) were modified to include *eGFP* (NC_011521.1 and NM_180996.1) (Stallings et al., 2002) and the recently identified cryptic exon of *Sry* (Miyawaki et al., 2020) (mkref function from cellranger). Protein coding genes and long non-coding RNAs were retained for further analysis. Gene-barcode matrices were generated with the cellranger count pipeline.

Seurat (version 2.3.4) R package and Scanpy (version 1.7.1) python package were used for downstream analysis (Butler et al., 2018; R Core Team (R Foundation for Statistical Computing), 2021; Wolf et al., 2018). Empty and low-quality barcodes were filtered out using a local minimum threshold on the UMIs *versus* barcode distribution of the raw matrices, as described in (Mayère et al., 2021b). The 12 datasets were merged (MergeSeurat function from Seurat), resulting in a combined dataset of 72,273 cells and 18,193 detected genes. Among the dataset, 39.2% (28,346) were from XX and 60.8% (43,927) from XY embryos, with 18.1% (13,060) from E9.0 samples, 20.7% (14,962) from E10.5, 23.1% (16,670) from E11.5 and 38.2% (27,581) from E12.5 (**Fig. S1C,D**). Raw UMI counts were normalized for library size (normalize_per_cell function from Scanpy), log-transformed (log1p function from Scanpy) and normalized for sequencing depth (regress_out function on UMI count from Scanpy).

Genes detected in more than 50 cells (14,394 genes) were used to compute independent component analysis (ICA) using their normalized value (RunICA function from Seurat, nics=100). Batch effect between replicates was corrected by computing the neighbourhood graph with the BBKNN python package (version 1.4.0) (neighbors_within_batch=3 and n_pcs=100) (Polański et al., 2020). For visualization, the corrected neighborhood graph was embedded in two dimensions using UMAP (Becht et al., 2019) (umap function from Scanpy with default parameters), and used for cell clustering with the Leiden algorithm (Traag et al., 2019) (leiden function from Scanpy, resolution=2.2).

To study AGP separation, we selected the 5,128 cells belonging to clusters 7 (CE adrenogonadal primordium), 17 (CE gonadal primordium) and 27 (adrenal primordium). We computed a force-directed graph (draw_graph function from Scanpy with default parameters) using previously computed corrected neighbour graph and ICA. A new clustering on this dataset was performed as described above using a resolution of 0.7.

For lineage reconstruction, connectivities between clusters were estimated (paga function from Scanpy with default parameters) (Wolf et al., 2019) and pseudotime calculated (dpt function from Scanpy with default parameters and root cell assigned from subcluster 1). Cells were classified according to their assigned path (adrenal: subclusters 1, 3, 6, 2, 5; gonadal: subclusters 1, 3, 6, 4, 0) and ordered according to their pseudotime score. Genes differentially expressed between adrenal and gonadal lineages, and between subclusters in pairs, were computed with a MAST test (FindMarkers function from Seurat), resulting in 1,737 differentially expressed genes. Genes were grouped according to their expression similarities, allowing the identification of specific expression patterns along pseudotime by hierarchical clustering using spearman distance (k-means=22, method=Ward.D2). Gene ontology enrichment analysis of biological processes was performed on each gene profile using ClusterProfiler (v4.1). Gene groups have been manually ordered to improve the visualization of the sequentially enriched profiles in either path.

For lineage reconstruction of the mesodermal derivatives, we removed erythroblasts/erythrocytes (clusters 44, 45, 49), hematopoietic progenitors (cluster 46), germ cells (clusters 5, 18, 25, 48), neural/sympathoadrenal cell types (clusters 28, 29, 31, 34, 41, 42, 47) and notochord (cluster 51), resulting in a dataset of 59,619 cells. Connectivities between clusters were estimated as described above using PAGA.

### Transcriptomic atlas annotation

Differential expression analysis on the 52 clusters for the complete dataset, or on the 8 subclusters of the subset, was calculated with a Mann-Whitney-Wilcoxon test (FindAllMarkers function from Seurat). Cluster-specific or -enriched genes were cross referenced with published cell type markers (highlighted in yellow in **Table S1**). Cell type annotation was done based on marker gene expression, developmental age, sex and sample type.

### Data and code availability

Raw data are available from the Gene Expression Omnibus (GSE156176). All code used for analysis is available upon request.

### Web interface

We provide an interactive web portal to explore our annotated single-cell adrenogonadal atlas (http://lmedapp731.unige.ch:5006/). This provides researchers the possibility to query for their genes of interest per cell types, age, and sex.

## Supplemental information titles and legends

**Figure S1 (related to Figure 1). Single-cell RNAseq metrics.**

**A, B.** Violin plots showing the number of UMIs, genes, UMIs/gene and mitochondrial gene proportion detected per cell for each condition (A) and for each cluster (B). Medians are indicated by a black line, and median of the entire dataset is indicated in bracket on the y axis. **C, D.** Bar plot showing cell number in each cluster, colored by age and sample type (C) and by sex (D). Corresponding pie chart showing cell percentage of the entire dataset is displayed on the right.

**Figure S2 (related to Figure 1). Single-cell transcriptomic atlas annotation.**

UMAP visualization of all cells colored by clusters of each major lineage (top) and expression level of representative markers (bottom).

**Figure S3 (related to Figure 3). Gene expression during adrenal and gonadal fate specification.**

Expression levels of representative genes related to Wnt and Rho signaling pathways (**A**), LPM-derived cardiac progenitor markers (**B**) and *HoxB* members (**C**) in a force-directed layout (top panel) and along pseudotime (bottom panel. x-axis: pseudotime; y-axis: normalized expression).

**Table S1 (related to Figure 1). Differentially expressed genes between clusters of the entire dataset.**

Reference markers genes are highlighted in yellow.

**Table S2 (related to Figure 2). Differentially expressed genes between subclusters of the AGP-AP-GP subset.**

**Table S3 (related to Figure 3). List of genes associated with the 22 profiles along pseudotime.**

**Table S4 (related to Figure 3). Gene ontology terms associated with the 22 profiles along pseudotime.**

**Note S1 (related to Figure 1). Additional transcriptomic atlas description.**

