## Supplementary material for "Single-cell transcriptomic profiling redefines the origin and specification of early adrenogonadal progenitors": Note S1

**Note S1 (related to Figure 1). Additional transcriptomic atlas description.**

The hematopoietic/endothelial lineage contains *Klf1*$^+$ erythroblasts (C45), *Hbb-bh1*$^+$/*Hba-a2*$^+$ primitive and *Hbb-bh1*$^-$/*Hba-a2*$^+$ definitive erythrocytes (C44 and C49, respectively), *Ptprc*$^+$/*Runx1*$^+$ hematopoietic progenitors (C46) and *Cdh5*$^+$ endothelial cells (C23).

Germ cells were identified by *Pou5f1* expression and consisted of several clusters, one of which contained primordial germ cells of both sexes (C25). C18 and C5 contained sex-enriched E12.5 cells, corresponding to *Lefty1*$^+$ prospermatogonia and *Stra8*$^+$ oogonia, respectively. Another germ cell population with higher mitochondrial gene expression clustered separately (C48, **Fig. S1B**), and could correspond to the nurse/dying cells described in mouse fetal ovaries by (Niu and Spradling, 2020). Cells of these four clusters are part of the dataset previously analyzed in (Mayère, et al., 2021).

The neural/sympathoadrenal lineage contains E9.0 *Foxb1*$^+$ neuroepithelial cells (C29), E9.0 *Plp1*$^+$ neural crest cells (NCC)/schwann cell precursors (SCP) (C42), and *Plp1*$^+$ NCC/SCP expressing both sympathetic progenitor marker (*Ret*) and SCP-to-chromaffin cell differentiation marker (*Htr3a*) (C34) (Furlan et al., 2017). We named this cluster "committed NCC/SCP". C34 forms two branches on the UMAP. The first is composed of cells at E10.5 that express a higher level of *Ret* and connect to early *Slc18a3*$^{low}$/*Chga*$^{low}$ sympathoblasts (C47). The second is composed of adrenal cells at E12.5 that express a higher level of *Hrt3a* and form a continuum with early *Slc18a3*$^-$/*Chga*$^{high}$ and late *Slc18a3*$^-$/*Chga*$^{high}$/*Foxq1*$^+$ chromaffin cells (C28 and C31, respectively). This is in agreement with the work of (Furlan et al., 2017) which revealed unexpected lineage segregation between sympathetic and adrenergic lineages at early stages. Late *Slc18a3*$^{high}$/*Chga*$^{low}$ sympathoblasts (C41) were obtained from adrenal gland samples at E12.5.

The smaller cluster of our dataset is composed of notochord cells at E9.0 cells that express the *Noggin* and *T* genes (C51).

Paraxial mesoderm-derived cells are divided into *Meox1*$^+$/*Pax3*$^+$ dermomyotome (C2), *Meox1*$^+$/*Pax9*$^+$ sclerotome (C12 and C19), *Myog*$^+$ myotome (C50) and *Acta2*$^+$ mural cells (C43) (Wasteson et al., 2008). An additional *Six1*$^+$/*Meox1*$^+$ cell population, negative for dermomyotomal, sclerotomal or myotomal markers is present and has been termed "paraxial mesoderm" (C24). All populations derived from the paraxial mesoderm, with the exception of mural cells, were derived from E9.0 samples.

The nephrogenic lineage was identified by *Pax2*$^+$/*Pax8*$^+$ expression, and includes E9.0 intermediate mesoderm (C26), *Tfap2b*$^{low}$/*Calb1*$^{high}$ caudal and *Tfap2b*$^{high}$/*Calb1*$^{low}$ rostral nephric duct (C33 and C37, respectively) (Sanchez-Ferras et al., 2021), *Cdh1*$^+$/*Jag1*$^+$ mesonephric tubules (C38) and *Cdh1*$^-$ mesonephric/metanephric mesenchyme (C21). C21 contains two subgroups of cells with mutually exclusive expression of the nephrogenic mesenchymal markers *Six2/Eya1* and mesonephric tubules markers *Lhx1/Osr2*, indicating that both uncommitted and committed tubular progenitors are present in this cluster.

In addition to the nephrogenic structures, the mesonephric ridges contain additional cell populations with, in particular, mesenchymal cells possessing mutually exclusive expression of the stromal markers *Foxd1/Cxcl12* and mesonephric mesenchymal markers *Wnt4/Gata2/Tbx3*. The latter group of cells, however, was distinct from *Wnt4*$^+$ mesonephric/metanephric mesenchymal cells (*i.e.* C21). We therefore classified these populations into the "mesonephros-associated cell types" lineage. This group includes the *Wnt4*$^+$ mesenchyme at E11.5 (C4), three clusters composed of *Wnt4*$^+$ and *Foxd1*$^+$ cells of different developmental stages (C0 for E10.5, C8 for E10.5/E11.5 and C1 for E11.5), and coelomic epithelial cells (*Upk3b*$^+$/*Krt7*$^+$/*Wnt4*$^+$) expressing markers of Müllerian duct progenitors *(Dach2/Lhx1/Pax8)* (C13), present as early as E10.5.

Lateral plate mesoderm lineage originated from E9.0 samples and contains *Foxf1*$^+$/*Tbx5*$^-$ splanchnic and *Foxf1*$^-$/*Tbx5*$^+$ somatic compartments (C32 and C39, respectively).

Overall, our dataset provides a comprehensive overview of the expected adrenogonadal cell types as well as surrounding tissues, present before and during AGP specification, as well as the more differentiated adrenogonadal lineages. This adrenogonadal developmental atlas is freely accessible through an interactive web portal (http://lmedapp731.unige.ch:5006/), allowing to query for genes of interest per cell types, age, sex and organ.
