## Supplementary Figures for "Single-cell transcriptomic profiling redefines the origin and specification of early adrenogonadal progenitors"

**Figure S1 (related to Figure 1). Single-cell RNAseq metrics.**

**A, B.** Violin plots showing number of UMIs, genes, UMIs/gene and mitochondrial gene proportion detected per cell for each condition (A) and for each cluster (B). Medians are indicated by a black line, and median of the entire dataset is indicated in bracket on the y axis. **C, D.** Bar plot showing cell number in each cluster, colored by age and sample type (C) and by sex (D). Corresponding pie chart showing cell percentage of the entire dataset is displayed on the right.

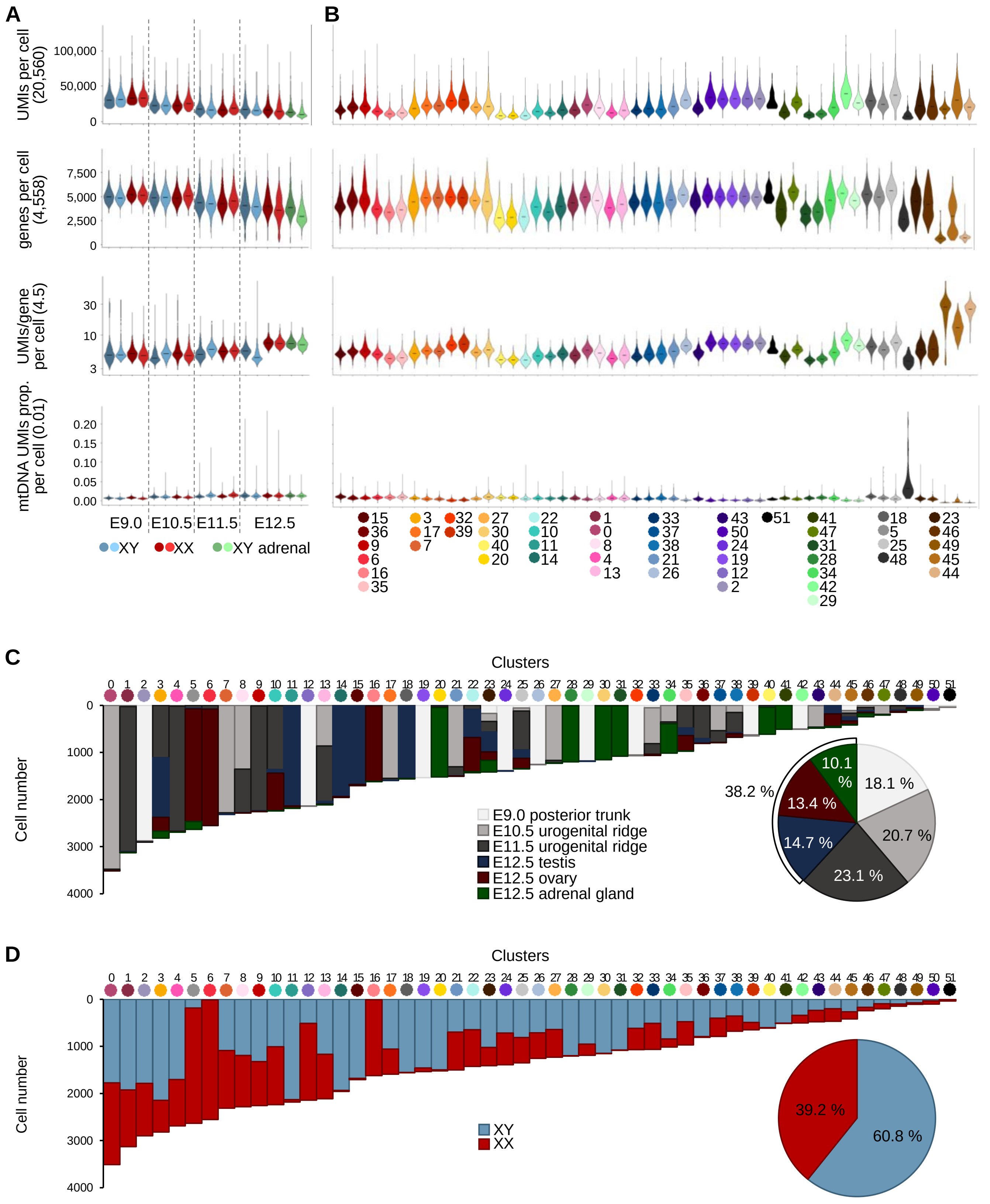

**Figure S2 (related to Figure 1). Single-cell transcriptomic atlas annotation.**
UMAP visualization of all cells colored by clusters of each major lineage (top) and expression level of representative markers (bottom).

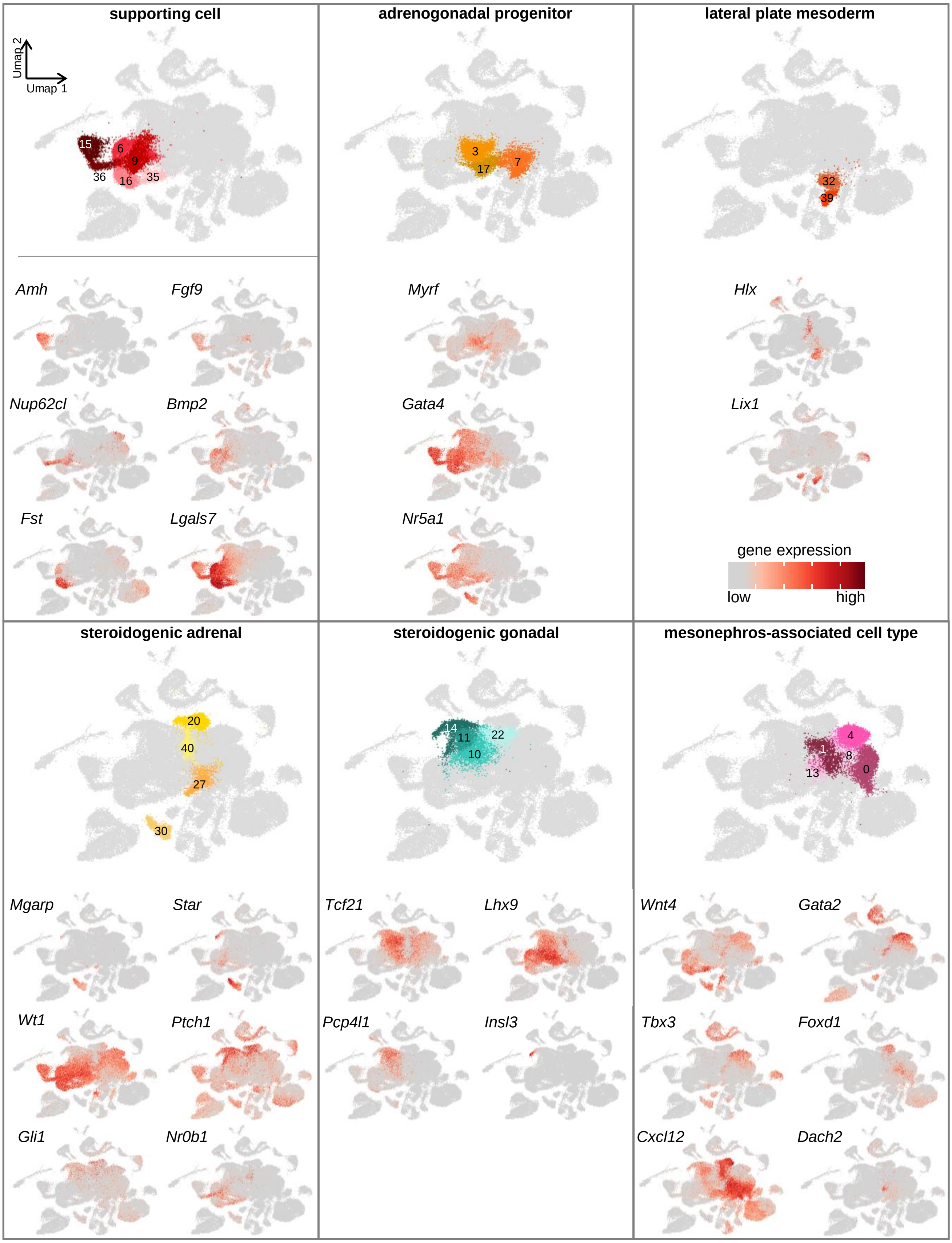

**Figure S2 (related to Figure 1) *continued*. Single-cell transcriptomic atlas annotation.**
UMAP visualization of all cells colored by clusters of each major lineage (top) and expression level of representative markers (bottom).

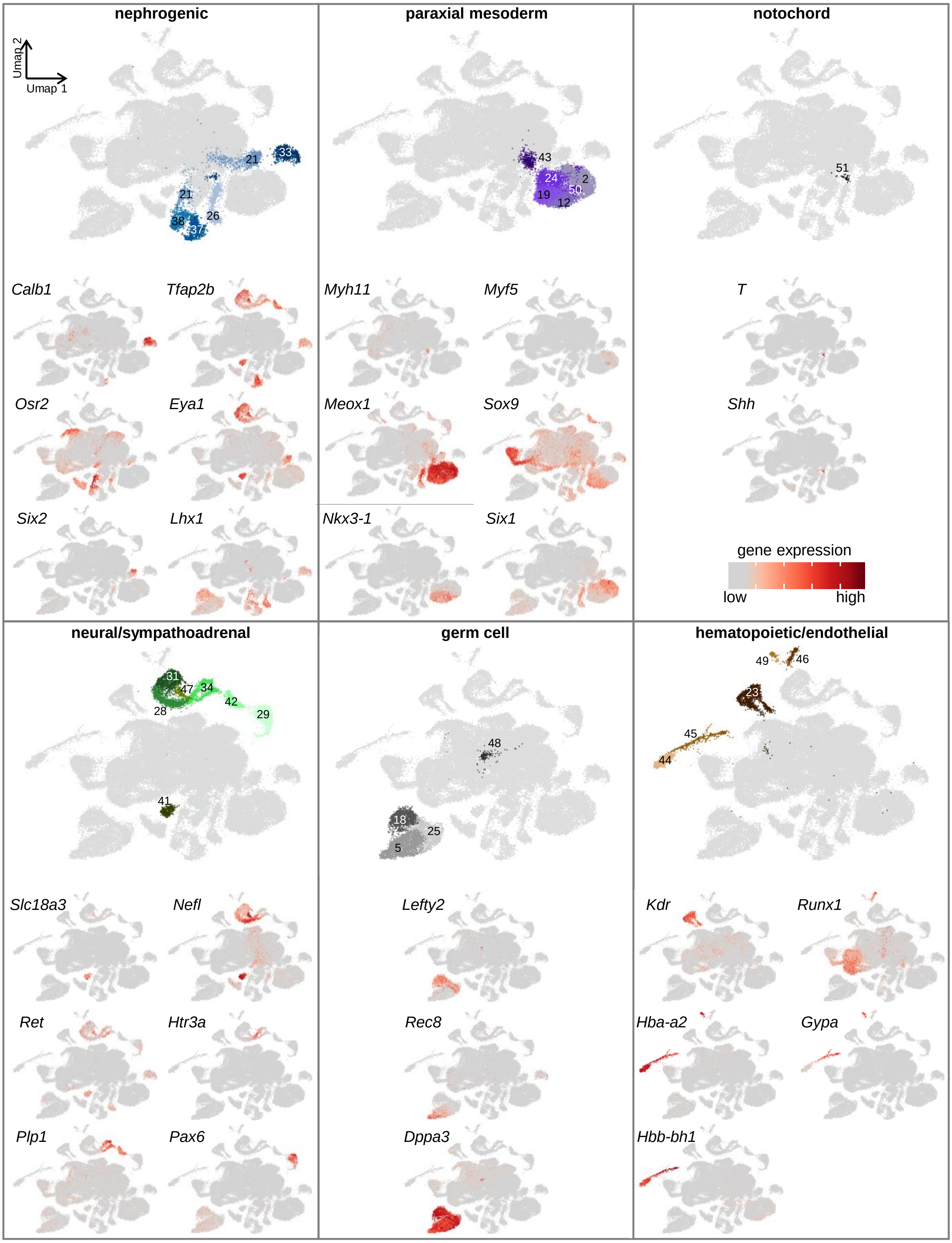

**Figure S3 (related to Figure 3). Gene expression during adrenal and gonadal fate specification.**
Representative genes expression level related to Wnt and Rho signaling pathways (**A**), LPM-derived cardiac progenitors markers (**B**) and *HoxB* members (**C**) in a force-directed layout (top pannel) and along pseudotime (bottom panel. x-axis: pseudotime; y-axis: normalized expression)**.**

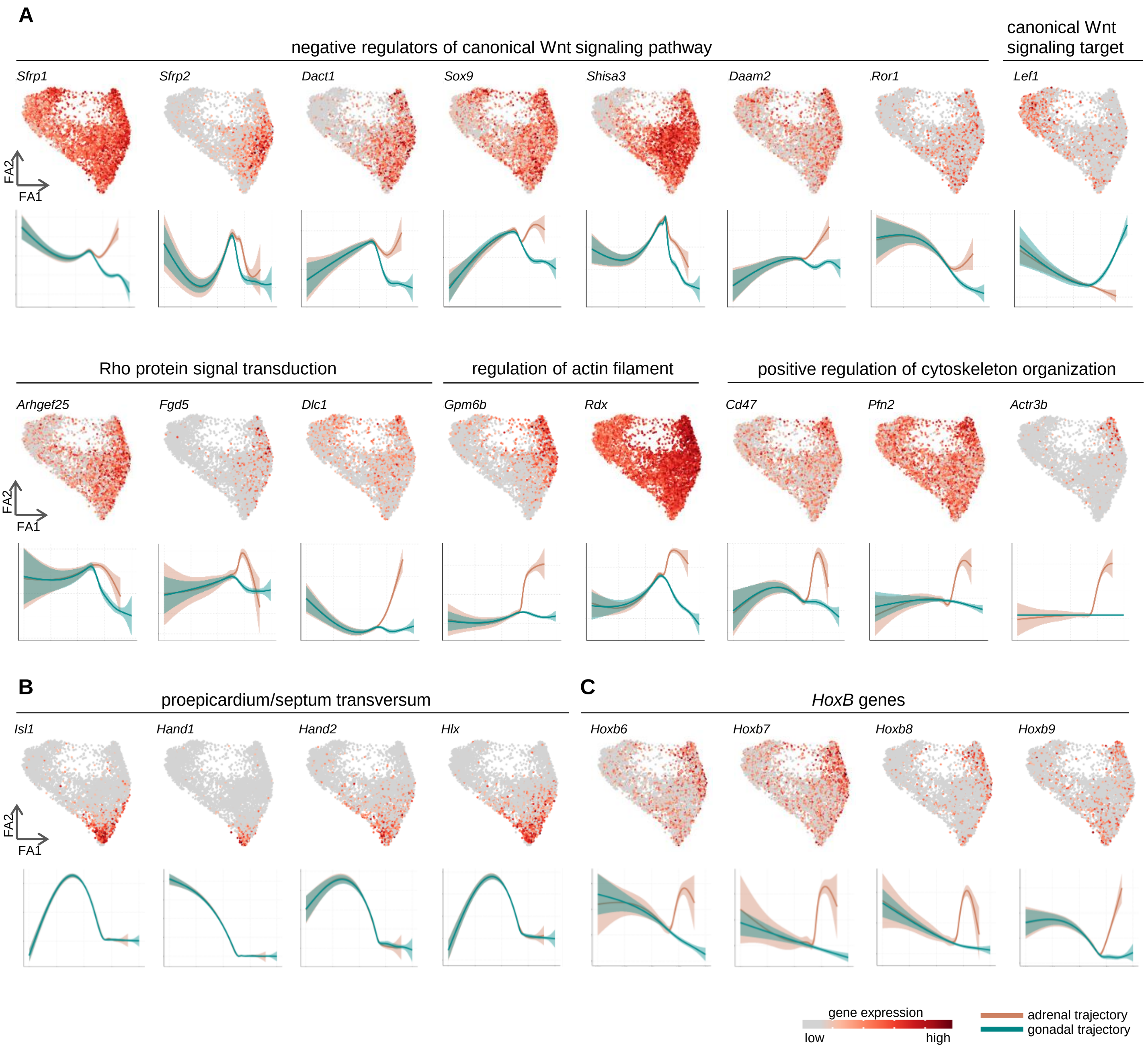
